# An Approximate Bayesian Computation Approach to Examining the Phylogenetic Relationships among the Four Gibbon Genera using Whole Genome Sequence Data

**DOI:** 10.1101/009498

**Authors:** Krishna R. Veeramah, August E. Woerner, Laurel Johnstone, Ivo Gut, Marta Gut, Tomas Marques-Bonet, Lucia Carbone, Jeff D. Wall, Michael F. Hammer

**Affiliations:** Arizona Research Laboratories Division of Biotechnology, University of Arizona, Tucson, AZ, USA; Department of Ecology and Evolution, Stony Brook University, Stony Brook, NY, USA; CNAG (Centro Nacional de Analisis Genomico), Baldiri Reixac 4, 08028 Barcelona, Spain; ICREA at Insitit de Biologia Evolutiva (CSIC/UPF), Dr. Aiguader 88, 08003 Barcelona, Spain; Department of Behavioral Neuroscience, Oregon Health and Science University, Portland, OR, USA; Institute for Human Genetics, University of California San Francisco, San Francisco, CA, USA

## Abstract

Gibbons are believed to have diverged from the larger great apes ∼16.8 Mya and today reside in the rainforests of Southeast Asia. Based on their diploid chromosome number, the family *Hylobatidae* is divided into four genera, *Nomascus*, *Symphalangus*, *Hoolock* and *Hylobates*. Genetic studies attempting to elucidate the phylogenetic relationships among gibbons using karyotypes, mtDNA, the Y chromosome, and short autosomal sequences have been inconclusive. To examine the relationships among gibbon genera in more depth, we performed 2nd generation whole genome sequencing to a mean of ∼15X coverage in two individuals from each genus. We developed a coalescent-based Approximate Bayesian Computation method incorporating a model of sequencing error generated by high coverage exome validation to infer the branching order, divergence times, and effective population sizes of gibbon taxa. Although *Hoolock* and *Symphalangus* are likely sister taxa, we could not confidently resolve a single bifurcating tree despite the large amount of data analyzed. Our combined results support the hypothesis that all four gibbon genera diverged at approximately the same time. Assuming an autosomal mutation rate of 1×10^−9^/site/year this speciation process occurred ∼5 Mya during a period in the Early Pliocene characterized by climatic shifts and fragmentation of the Sunda shelf forests. Whole genome sequencing of additional individuals will be vital for inferring the extent of gene flow among species after the separation of the gibbon genera.

## Introduction

The family *Hylobatidae*, commonly known as gibbons, represents the most divergent lineage of the ape superfamily, separating from the lineage leading to the great apes ∼16.8 Mya (Carbone et al. in review.). Sometimes known as small apes, gibbons demonstrate substantial morphological differentiation from the great apes; their much smaller bodies are highly adapted to an arboreal mode of locomotion in the rainforests of Southeast Asia. They also demonstrate very little sexual dimorphism that may, in part, be related to their generally monogamous mating patterns (FUENTES 2000) (although some gibbon species develop differences in coat color at sexual maturity).

Each species demonstrates distinct ‘call’ and ‘song’ types (GEISSMANN 2002); however, attempts to classify gibbon species and genera based solely on morphological features have been problematic (FUENTES 2000; MOOTNICK 2006). Primarily on the basis of their karyotypes, gibbons are now divided into four major genera, with *Nomascus*, *Symphalangus*, *Hylobates* and *Hoolock* each possessing 52, 50, 44, and 38 diploid chromosomes, respectively. While many genetic studies have been performed, including a number based on karyotypes (GEISSMANN 2002; MÜLLER *et al.* 2003), mitochondrial DNA (HAYASHI *et al.* 1995; TAKACS *et al.* 2005; MONDA *et al.* 2007; WHITTAKER *et al.* 2007; VAN NGOC *et al.* 2010; MATSUDAIRA and ISHIDA 2010), Y chromosomes (CHAN *et al.* 2012), ALU repeats (GEISSMANN 2002; MEYER *et al.* 2012), and short stretches of autosomal sequence (FUENTES 2000; MOOTNICK 2006; KIM *et al.* 2011; WALL *et al.* 2013), the phylogenetic relationships among the four gibbon genera remain unresolved, with at least seven different topologies being supported by different data.

A recent study examined ∼1.5 Mb of orthologous autosomal sequence generated by 2nd generation sequencing from one individual representing each of the four genera (GEISSMANN 2002; MÜLLER *et al.* 2003; WALL *et al.* 2013). This study, too, was inconclusive and suggested that the gibbon genealogy demonstrates substantial incomplete lineage sorting (ILS). However, the experimental design was limited by the lack of a suitable reference genome (short reads were aligned to highly divergent human hg19 assembly). To examine the species tree relationships among gibbons, as well as estimate key demographic parameters such as the time when the various gibbon genera diverged, we generate whole genome sequence data from eight individuals representing all four gibbon genera and utilize the newly released gibbon (nomLeu1) reference genome (Carbone et al. in review) for mapping and variant calling. Then we apply a novel coalescent-based Approximate Bayesian Computation (ABC) approach that can handle large amounts of sequence data and that corrects for potential sequencing error and reference genome mapping bias.

## Materials and Methods

Blood and tissues were obtained in agreement with protocols reviewed and approved by the Gibbon Conservation Center. More details on all aspects of the methods are provided in the Supplementary Information.

### Sequence Generation

DNA was extracted from blood or cell lines, and paired-end libraries were prepared with the Illumina TruSeq chemistry. Libraries were sequenced on the HiSeq 2000 platform, generating 2×100 bp reads. Multiple runs were performed to generate a minimum of 10X mean coverage on each sample after all post-processing. Mean coverage ranged from 11.5X to 19.5X.

### Read Mapping and Variant Calling

Trimmed reads were aligned to nomLeu1 with Stampy (v. 1.0.17) (HAYASHI *et al.* 1995; TAKACS *et al.* 2005; MONDA *et al.* 2007; WHITTAKER *et al.* 2007; VAN NGOC *et al.* 2010; MATSUDAIRA and ISHIDA 2010; LUNTER and GOODSON 2011). For the two *N. leucogenys* (NLE) samples, Stampy was used in its “hybrid mode” where alignment with BWA (v. 0.5.9) (LI and DURBIN 2009; CHAN *et al.* 2012) is attempted first. A substitution rate of 0.001 was specified, along with BWA minimum seed length of 2, fraction of missing alignments 0.0001, and quality threshold 10. For the non-NLE samples, Stampy was used with a substitution rate of 0.015 (KIM *et al.* 2011). Local realignment at indel sites was performed with the Genome Analysis Toolkit (GATK, v. 1.4-37) (MCKENNA *et al.* 2010; DEPRISTO *et al.* 2011). PCR duplicates were removed with samtools. GATK UnifiedGenotyper was run separately on the two samples from each genus and Single Nucleotide Variants (SNVs) and indels with a quality score of at least 50 were retained to create a mask of variant sites to be excluded from base quality score recalibration. The GATK indel realignment tool was run again to standardize alignment of indels across all samples. UnifiedGenotyper from GATK version 2.1-11 (to allow multiallelic calling) was used to produce a final set of SNVs and indels. Each site was annotated with the consensus quality score of the nomLeu1 reference sequence.

### Masks

The nomLeu1 genome is composed of 17,968 contigs, ranging in size from 2,496 bases to ∼74 MB. As small loci may be compressed, and represent duplications in the gibbon genome that have not been properly separated during the assembly process, we masked out all scaffolds less than 1MB in length, yielding 273 scaffolds that span ∼2.73 GB. UCSC's gibbon-human pairwise alignments where used to identify non-autosomal sequence. Specifically, gibbon loci that aligned to human X, Y or M in UCSC's “net” alignments (KENT *et al.* 2003) were masked, along with locations in the gibbon genome that were not primary alignments to locations in the human genome. Further, locations where the gibbon reference quality was below a phred-quality of 50, repeats (identified by Tandem Repeat Finder (BENSON 1999) or by RepeatMasker (SMIT *et al.* 1996)), LAVA elements identified in Carbone et al.(in review), CNVs with an estimated ploidy >2.5 in any sample (also identified in Carbone et al.(in review), infinite sites violations, positions where any sample has less than 7x coverage, or more than their 95^th^ percentile read depth, and bases within 3bp of any indel called were excluded, unless otherwise specified, from downstream analysis.

### Exome Validation and Calibration

Exome capture using the TruSeq Exome Enrichment Kit (Illumina) was performed on one NLE sample (Vok, 116x coverage) and one SSY sample (Monty, 64x coverage), and the resulting data were run through the pipeline described above. Our WES exome calibration makes the following simplifying assumptions: (1) after masking, any exome base with 30x <= coverage <= 200x is considered “correct” (called without error), (2) read-depth and mapping bias (whether or not the sample belongs to the same taxon as the reference) account for all genotyping errors observed, and (3) all false negatives (i.e., SNPs present in the exome, but not present in the whole genome data) are singletons. We separate errors, *E*, into two categories; errors, *S*, involving singleton polymorphisms (defined with respect to the nonLeu1 reference), and genotyping errors when the polymorphism is segregating with a non-reference allele present in two or more chromosomes. For the former, we concern ourselves with the rate of singleton calling per sample, i.e., the fraction of the singletons in our whole genome data that are called in the exome capture data. For the latter we create confusion matrices *M* over the set of genotype calls {Reference, Heterozygous, Alternative} to describe the type and probability of all possible errors. This results in an error function *E* = <*S*, *M*> which transforms perfectly correct data into data reflective of the error processes that are likely to have occurred during whole genome sequencing and post processing. In order to apply our error correction to samples that did not undergo exome sequencing, our final version of *E* is based on an empirical distribution of read depths for each sample, and whether or not that sample is from the same genus as the reference.

### Machine learning for SNP-based analysis

The machine learning (ML) program Weka 3.6.8 (HALL *et al.* 2009) was used to classify the whole genome genotype data at all called segregating sites regardless of quality, with the aim of finding a subset of very high quality sites for use in our PCA and calculation of F_ST_. Using the same definition of “correct” as above, we generated a training set of all sites that were incorrectly called in the genome, and a random and equally sized sampling of sites that were called correctly for both our NLE and our non-NLE (SSY) sample. A variety of features from the GATK output as well as whether the call is from the NLE or the non-NLE sample, and the combined p-value of the distribution of read depths observed at the site were used in the machine learning analysis. Four ML algorithms– multilayer perceptron, ridor, rotation forest and classification by regression– showed reasonable performance (75%-85% accuracy). After various optimization procedures, we classified a genotype call as correct if all four classifiers predicted that the genotype was correct, and we classified a site as correct if all genotypes at a site were classified as correct. PCA was performed using smartpca (PATTERSON *et al.* 2006) and visualized using R.

### ABC analysis

ABC analysis was performed on two data sets containing independent loci of small enough length such that intra and interlocus recombination could be ignored in our simulations. Set 1 included 12,413 non-genic loci consisting of 1 kb of total callable sequence across a contiguous stretch of no more than 3 kb separated by at least 50 kb and at least 50 kb from the nearest exon. Set 2 included 11,323 genic loci consisting of 200 bp of total callable sequence across a contiguous stretch of no more than 4 kb separated by at least 1 kb (this distance will likely violate our assumption of independence but increasing this distance substantially decreased the number of usable loci and thus reduced our power to a greater extent), with an allowance of a maximum of 100 bp of the locus lying adjacent to an exon and the rest lying in the exon (**Fig S1**). In addition to the masks and coverage filters described above, we also masked CpG consistent sites as well as conserved phastCons (SIEPEL *et al.* 2005) elements inferred from primate genomes with a further 100 bp padding either side of the element. Variant sites were polarized against the aligned human reference genome, hg19.

To account for mutation rate heterogeneity among loci we estimated relative sequence divergence for all loci, taking the average sequence divergence for each of the eight gibbon individuals from hg19. These individual locus estimates were then normalized around a mean of 1, allowing us to follow the approach of Rannala and Yang (RANNALA and YANG 2003) and scale θ for each individual locus in our demographic simulations. We computed the following summary statistics to describe the data for every pair of populations across all loci: mean number of shared derived polymorphisms, mean number of private derived polymorphisms in each population and the mean number of private fixed sites in each population

We treated all possible phylogenetic relationships among the four gibbon genera as distinct models. The models are described by two classes of parameters, mean population nucleotide diversity, θ, and branch lengths in units of expected number of substitutions, τ (thus mutation rates per site per generation do not need to be explicitly stated during the analysis). Priors ranged between 0.0001–0.03 for all θ and τ parameters. Unless stated all prior distributions for all demographic parameters are all uniformly distributed on a log_10_ (x) scale. Simulations were performed using a version of ms (HUDSON 2002) modified for Python that allowed fast parallel processing. Error models (*S_i_*, *M_i_*) from the exome validation were generated specifically for the coverage in each individual at the specific regions considered (i.e. at the non-genic and genic loci) and incorporated into the simulations by randomly dropping singleton heterozygous sites at rate *S_i_* and assigning a genotype at segregating sites with two or more derived sites based on *M_i_*. When estimating model parameters we utilized ABCtoolbox (WEGMANN *et al.* 2010), which implements a general linear model (GLM) adjustment (LEUENBERGER and WEGMANN 2010) on retained simulations. Before ABC analysis the full set of summary statistics was transformed into (PLS) components (WEGMANN *et al.* 2009) and we used the change in (RMSE) to guide the choice of number of components. We used the logistic regression (LR) method previously described (FAGUNDES *et al.* 2007) to perform model choice. 1% of simulations were retained for the GLM (parameter estimation) and LR (model choice) adjustments.

2% ancestral state misidentification was incorporated into simulations by calculating the expected number of sites likely to experience a mutation along the hg19 lineage for each loci (1000 bp x 2% = 20 sites). The number of sites to actually “flip” (i.e. assign the wrong ancestral state) for each loci during a simulation is drawn from a Poisson distribution with this mean. These sites are then randomly assigned to a position along the locus, though only positions that are found to segregate amongst the gibbon chromosomes need to be flipped.

### G-PhoCS analysis

The Markov Chain Monte Carlo (MCMC) Bayesian coalescent-based method described by Gronau et al. (GRONAU *et al.* 2011) was performed using the software G-PhoCS to estimate θ and τ values for a bifurcating tree (we ignored the effect of migration). On this occasion we included a human haploid sequence (hg19) as an outgroup for the overall gibbon phylogeny (rather than just to infer the ancestral state as in the ABC analysis). The same 12,431 1 kb loci and assumed best bifurcating species tree from the ABC analysis described above were utilized and the mutation rate was fixed individually for each loci as above using the normalized divergence values. The gamma prior for θ was set to be relatively broad and uninformative and the same for all present and ancestral populations with α = 2 and β =1,000. Gamma priors for τ were also set to be relatively uninformative, with the α value always 2. However, either *a*) β was set as 200 for all τ values or *b*) individual β values were set for each τ such that the mean value reflected rough estimates from the ABC analysis or for the human/gibbon split time from Carbone et al.(in review) (**Table S1**). We ran three independent MCMC chains for both prior settings *a*) and *b*). We allowed 10,000 samples as burn-in followed by 100,000 samples for estimating parameters. All parameters converged much quicker than the utilized burn-in period, and all six runs converged to the same parameter space. Results were processed using the software Tracer (http://tree.bio.ed.ac.uk/software/tracer/).

## Results

### 2^nd^ generation sequencing and validation

We performed 2^nd^ generation whole genome sequencing (WGS) on two individuals (one male and one female) from each of the four gibbon genera (**Table 1**). For our *Nomascus* samples, represented by the species *leucogenys* (NLE, the northern white-cheeked gibbon), the two individuals examined differed from the (NCBI Project 13975 GCA_000146795.1) nomLeu1 reference genome. For our *Hylobates* samples (the most diverse genus with ∼13 species), we examined one individual each from the *H. moloch* (HMO, Javan gibbon) and *H. pileatus* (HPI, Pileated gibbon). Our *Symphalangus* sample is represented by two individuals from the species *syndactylus* (SSY, Siamang gibbon). It is important to point out that the two *Hoolock* samples from the *leuconedys* species (HLE, Eastern hoolock gibbon) represent the only wild born individuals present in the study, whereas all other individuals were captive-born (i.e., offspring of individuals living in zoos). We also mention that matings between different gibbon species (and even different genera) are known to result in viable offspring in captivity (MYERS and SHAFER 1979; MOOTNICK 2006; HIRAI *et al.* 2007). If any of the individuals in our sample are indeed hybrids between different species, our analysis may be affected in unexpected ways.

**Table 1:** Gibbon samples undergoing 2^nd^ generation sequencing.

| Chr<br>Nb | Genus | Species | Common<br>Name | Code | Sex | Origin | Mean<br>Coverage |
| --- | --- | --- | --- | --- | --- | --- | --- |
| 52 | *Nomascus* | *Nomascus* | Northern white- | NLE | M | parents WB | 13.78 |
|  |  | *leucogenys* | cheeked |  | F | parents WB | 11.50 |
| 50 | *Symphalangus* | *Symphalangus* | Siamang | SSY | M | sire WB, dam CB | 12.80 |
|  |  | *syndactylus* |  |  | F | parents CB | 19.53 |
| 38 | *Hoolock* | *Hoolock* | Eastern | HLE | M | WB | 19.15 |
|  |  | *leuconedys* | hoolock gibbon |  | F | WB | 14.36 |
| 44 | *Hylobates* | *Hylobates*<br>*pileatus* | Pileated gibbon | HPI | M | parents WB | 14.33 |
| 44 | *Hylobates* | *Hylobates*<br>*moloch* | Javan gibbon | HMO | F | sire WB, dam CB | 12.96 |

WB = wild born, CB = born in captivity

After post-processing the sequence data we obtained a mean coverage of 15X (min = 11.5X, max =19.5X) (**Fig S2**). As previous work has indicated a relatively high divergence between gibbon genera, we attempted to incorporate potential reference bias into our post-processing by utilizing a higher substitution rate (1.5%) when mapping sequence reads for non-NLE samples, and by using a hybrid mapper, Stampy (LUNTER and GOODSON 2011) to increase sensitivity. To validate our variant calling we performed high coverage whole exome capture sequencing (WES) on one NLE individual and one non-NLE sample (the male SSY sample). Mean coverage for WES data was 116X (compared with 14x for WGS data) and 64X (compared with 13X for WGS data), respectively. Human-based exome capture has been shown to be effective in primates as diverged from humans as macaques (JIN *et al.* 2012). Utilizing only exome calls with coverage between 30x and 200x we found slightly greater concordance between the WGS and WES data for the NLE (99.6%) *versus* non-NLE samples (99.4%) (**Table S2)**. Noticeably when only examining singleton variants, calling was markedly better in the reference taxa (∼99% of exome-called sites identified in the WGS data) than in the non-reference taxa (∼96%), suggesting reference biases may still exist in our data for rare variants in non-reference taxa.

### Genetic diversity among gibbon genera

Within genera diversity, assessed for this dataset by Carbone et al. (in review), demonstrated that NLE samples had the highest level of nucleotide diversity (π ∼2.2×10^−3^), while values as low as ∼7.3×10^−4^ were observed in the HPI sample. Nucleotide diversity for the HMO sample was also relatively high at ∼1.7×10^−3^, followed by SSY (∼1.4×10^−3^), and then the two wild born HLE (∼8×10^−3^). By way of comparison, π ranges from approximately 0.5–1.0×10^−3^ in humans, 1.8×10^−3^ in western lowland gorillas, and 2.3×10^−3^ in Sumatran orangutans (PRADO-MARTINEZ *et al.* 2013). To examine the relative levels of genetic differentiation among the gibbon genera we performed Principal Components Analysis (PCA) on the individual samples. For this analysis we examined di-allelic SNPs called in all individuals. High-quality SNPs were identified by using concordance with the WES data to train a machine-learning (ML) algorithm to predict highly confident SNPs across the whole genome and in samples that did not undergo WES. In addition, to ensure independence of SNPs, we randomly selected sites that were separated by at least 100 kb when on the same scaffold. This resulted in a dataset of 25,531 high quality genome-wide independent SNPs. The first four principal components accounted for 40.2%, 31.2%, 24.6% and 3.5% of the variation, respectively (**Fig S3A and B**). The four genera showed substantial genetic differentiation and were clearly separated in the PCA plot in the first two components, though no clear inter-genera phylogenetic relationship emerged. Individuals from the same species showed high similarity suggesting limited inter-genera hybridization or contamination. The two *Hylobates* species could be clearly distinguished in PC4. We were also able to reproduce the same patterns when only using a random subset of ∼200 SNPs (**Fig S3C and D**), suggesting it may be possible to perform relatively low coverage shotgun sequencing from a number of different gibbon species and use a similar approach to this in order to identify a small yet powerful set of species specific SNPs. This could be particularly important for management of gibbons in zoos when it can often be difficult to distinguish different species or even genera based on fur alone, often leading to accidental hybrids (TENAZA 1985).

### A Coalescent-based ABC Analysis of the Gibbon Phylogeny

Unless species branch lengths are several orders of magnitude larger than the expected time to the most recent common ancestor of sequences within a species, it is important to model stochasticity in the distribution of gene trees across loci when inferring an underlying species tree (ROSENBERG and NORDBORG 2002). Current Bayesian coalescent-based methods such as BEAST (DRUMMOND and RAMBAUT 2007) that explicitly take into account sequence and population divergence simultaneously to infer species trees are generally computationally intractable for large datasets (BRYANT *et al.* 2012). Therefore, in order to infer the species topology for gibbon genera we developed an Approximate Bayesian Computation (ABC) (BEAUMONT *et al.* 2002) method for inference of a species tree with four taxa that can handle large amounts of sequence data, is not dependent on haplotype phase, and incorporates information derived from our WES validation.

Analogous to the likelihood approach of Gronau et al.(GRONAU *et al.* 2011), the data required for this ABC method are short, independent loci as we assume no intra-locus recombination and free recombination between loci. The latter is a necessary convenience given that no recombination map is currently available for gibbons. Thus, we assembled a set of independent ‘non-genic’ sequences that mapped at least 50 kb away from genes (∼12,000 1 kb loci) and that excluded CpG consistent sites as well as evolutionarily conserved elements (SIEPEL *et al.* 2005) (**Fig S1**). Mutations detected in these loci are expected to represent neutral variation and to evolve at a relatively constant rate. To reduce reference-mapping bias, we also assembled an analogous set of independent ‘genic’ loci that span exons (∼11,000 200 bp loci) and that should have lower diversity, recognizing that these loci may have been subjected to natural selection, which may bias parameter estimates.

Analysis of simulated data demonstrated that our method had 88.4% power to detect the correct topology from randomly drawn datasets, with the correct model among the three highest posterior probabilities 99% of the time (**Fig. S4**). A more targeted power analysis demonstrated that the method is only likely to fail when an internal branch is extremely small (almost instantaneous in evolutionary terms) or when the total height of the tree is on the order of 0.001 (equivalent to about 1 million years) (**Fig S5**), which is unrealistic for gibbons. A more detailed discussion of the validation can be found in the **Supplementary Information**.

As most ABC analyses are based on performing simulations to approximate an otherwise intractable likelihood function, we were also able to incorporate into the simulations our whole WES validation findings by modeling sequence errors (missing singletons and incorrect genotype calls at other segregating sites) that occurred in the real data. A full description of how our WES validation was incorporated into this analysis is given in the **Supplementary Information**. Prior to the ABC analysis we examined the one-dimensional distribution for each individual summary statistic from 10,000 random error-corrected simulations and found a good fit to our non-genic and genic observed data, while a Principle Component Analysis also demonstrated a good multidimensional fit (**Fig S6**).

**Table 2** shows the posterior probabilities for the ABC analysis for all phylogenetic models using the corrected and uncorrected simulations for both the non-genic and genic loci. No topology dominates the analysis, with three to four topologies having posterior probabilities >10% in the corrected simulations. The best topology using non-genic and genic loci for the corrected simulations differ, and both still maintain relatively low posterior probabilities of 19%. Two topologies appear most prominent with posterior probabilities >10% in all four analyses and the highest means across all four analyses and both (genic and non genic) exome-corrected analyses. One is the most frequently observed topology in the sequence divergence analysis (((SSY, HLE), NLE),(HPI,HMO)) of Carbone et al. (in review) and the other is a related topology where (HPI, HMO) and NLE are swapped as the most external groups with HLE and SSY remaining as sister taxa. Together the posterior probability for both these related topologies sum to 30–32%. However, in general the posterior probabilities are lower than typically observed in our simulations suggesting that we have little confidence in the true topology. This is consistent with the hypothesis of a rapid radiation of gibbon species from a large ancestral population.

**Table 2:** Posterior probabilities for the 15 possible 4-population topologies for non-genic and genic loci

| Topology | Non-Genic |  | Genic |  |
| --- | --- | --- | --- | --- |
|  | Corrected | Uncorrected | Corrected | Uncorrected |
| **(( (SSY, HLE) NLE) (HPI, HMO) )** | 0.16 | 0.15 | 0.19 | 0.15 |
| (( ( (HPI, HMO) NLE) SSY) HLE) | 0.19 | 0.14 | 0.11 | 0.08 |
| (( (SSY, HLE) (HPI, HMO) ) NLE) | 0.14 | 0.23 | 0.13 | 0.19 |
| (( ( (HPI, HMO) NLE) HLE) SSY) | 0.13 | 0.11 | 0.06 | 0.05 |
| (( (NLE, HLE) SSY) (HPI, HMO) ) | 0.06 | 0.05 | 0.10 | 0.08 |
| (( ( (HPI, HMO) SSY) NLE) HLE) | 0.07 | 0.06 | 0.08 | 0.07 |
| (( ( (HPI, HMO) SSY) HLE) NLE) | 0.05 | 0.07 | 0.07 | 0.14 |
| (( (HPI, HMO) NLE) (SSY, HLE) ) | 0.05 | 0.04 | 0.05 | 0.03 |
| (( (NLE, SSY) HLE) (HPI, HMO) ) | 0.03 | 0.03 | 0.06 | 0.04 |
| (( (NLE, HLE) (HPI, HMO) ) SSY) | 0.04 | 0.03 | 0.04 | 0.04 |
| (( (NLE, SSY) (HPI, HMO) ) HLE) | 0.03 | 0.03 | 0.03 | 0.02 |
| (( ( (HPI, HMO) HLE) SSY) NLE) | 0.02 | 0.04 | 0.03 | 0.06 |
| (( ( (HPI, HMO) HLE) NLE) SSY) | 0.02 | 0.02 | 0.02 | 0.02 |
| (( (HPI, HMO) SSY) (NLE, HLE) ) | 0.01 | 0.01 | 0.03 | 0.02 |
| (( (HPI, HMO) HLE) (NLE, SSY) ) | 0.01 | 0.01 | 0.01 | 0.00 |

Bold type indicates the topology identified using sequence divergence in Carbone et al. (in prep)

### Estimation of Parameters describing Gibbon Demography

To estimate when this rapid radiation may have taken place we constructed a model where all four genera diverge simultaneously (an instantaneous radiation model), with a subsequent divergence of the two *Hylobates* species. This resulted in a model with seven θ and two τ (in units of expected number of mutations) parameters. The summary statistics from the non-genic loci were transformed into partial least squares (PLS) components to infer the demographic parameters. Posterior distributions and parameter estimates are shown in **Table S3** and **Fig. S7.** These results are based on 15 PLS components, the value at which the best reduction in the root mean square error (RMSE) was observed across all parameters, **Fig. S8**, and for which the 95% CIs for τ were relatively reliable based on 1,000 pseudo observed datasets.

Values of π described above were within the 95% CI for the θ values estimated by the ABC analysis for present-day species and showed the same relative pattern with the highest value in the NLE and lowest value in the HPI sample. The divergence time, τ_1_, for the two *Hylobates* samples was ∼50% less than that for the divergence time of the four gibbon genera, τ_2_, which is consistent with the relative difference in sequence divergence of ∼0.5% seen in Carbone et al. (in review.). Because the priors were log_10_ scaled, the associated 95% CI values potentially could be larger in absolute values (i.e. 10^^val^) than if the observed posterior distribution had been shifted towards a smaller branch length. Therefore, we re-ran the ABC analysis using un-scaled flat priors for the two τ values, which resulted in highly similar median values but much narrower 95% CIs (**Table 3, Table S4, Fig. S9**). We note that the CIs were somewhat anticonservative as assessed by simulated datasets (see column “HDPI 95% fit” of **Table 3** **and Table S4**). When we assume a µ of 1×10^−9^ per base per year * 3/4 (to take into account that we excluded CpG sites (HODGKINSON and EYRE-WALKER 2011)) this results in an estimate for the time of the gibbon radiation of 1.6+3.5 = 5.1 Mya (τ_1_−τ_2_ combined limits of 95% CI 2.5–7.7 Mya) and a split time of 1.6 Mya (95% CI 0.6–2.9 Mya) for the two *Hylobates* samples. In addition, assuming 10 years per generation for gibbons (HARVEY *et al.* 1987) and thus a µ of 7.5×10^−9^ per generation, N_e_ for extant species varies from 57,000 (NLE) to 7,500 (HPI). Interestingly, the ancestral gibbon N_e_ is estimated to be much larger at 132,000 (107,000–162,000) (**Fig. 1a**) as would be expected under a model of substantial ILS. It should be noted that the estimate of the ancestral *Hylobates* population size (based on θ_T1_) may be somewhat unreliable as the regressed posterior distribution shows a major shift from the raw retained posterior distribution (**Fig. S9)** and the RMSE analysis showed this was the θ value for which there was least power for inference using the smallest number of PLS components (**Fig. S10)**.

**Figure 1a:**
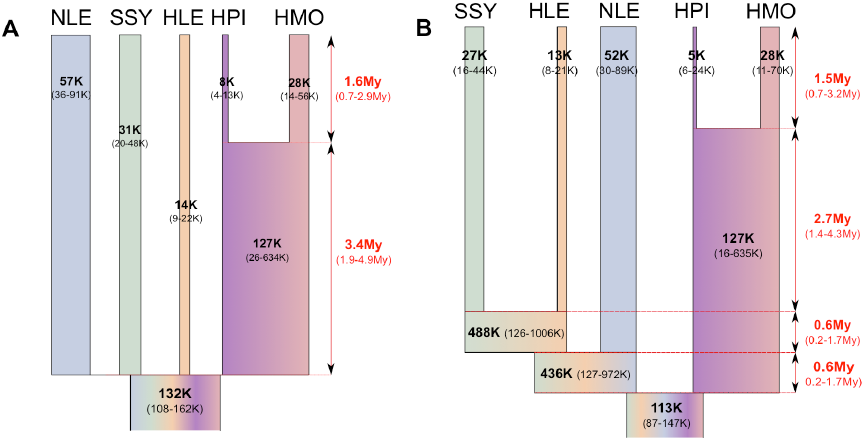
Parameter estimates for the instantaneous radiation model for gibbon genera. µ = 7.5×10^−9^ /site/generation, 10 years per generation.

**Table 3:** Posterior estimates for an instantaneous speciation model for gibbon genera using a flat prior for τ

| Parameter | HDPI 95% fit<sup>a</sup> | Posterior Estimation<sup>b</sup> |  |  |  |
| --- | --- | --- | --- | --- | --- |
|  |  | Mode | Median | HDPI 95 |  |
|  |  |  |  | Lower | Upper |
| $\theta_{NLE}$ | 0.930 | 1.71E-03 | 1.72E-03 | 1.07E-03 | 2.73E-03 |
| $\theta_{SSY}$ | 0.936 | 9.25E-04 | 9.24E-04 | 5.97E-04 | 1.43E-03 |
| $\theta_{HLE}$ | 0.937 | 4.17E-04 | 4.17E-04 | 2.63E-04 | 6.58E-04 |
| $\theta_{HPI}$ | 0.968 | 2.24E-04 | 2.25E-04 | 1.30E-04 | 3.92E-04 |
| $\theta_{HMO}$ | 0.974 | 8.29E-04 | 8.32E-04 | 4.13E-04 | 1.68E-03 |
| $\theta_{T1}$ | 0.958 | 3.54E-03 | 3.80E-03 | 7.69E-04 | 1.90E-02 |
| $\theta_{Tanc}$ | 0.964 | 3.97E-03 | 3.97E-03 | 3.23E-03 | 4.86E-03 |
| $\tau_1$ | 0.905 | 1.05E-03 | 1.23E-03 | 5.01E-04 | 2.18E-03 |
| $\tau_2$ | 0.911 | 2.69E-03 | 2.59E-03 | 1.41E-03 | 3.63E-03 |

<sup>a</sup>A metric demonstrating how often known simulated values (n=1,000) fell within the calculated 95% CI, which gives a guide to the reliability of these CI's for real data. <sup>b</sup>Calculated using 15PLS components, 1,000,000 simulations and retaining 1%. All priors ranged from 0.0001-0.03 when $\log_{10}$ scaled.

One potential source of error in estimating parameters is ancestral state misidentification due to back mutations along the human lineage, which was used as an outgroup (HERNANDEZ *et al.* 2007). Our simulated data assumed an infinites sites model. Assuming a human-gibbon split time of 16.8 Mya and µ of 1×10^−9^ per base per year, each site has ∼98% chance ((1–1×10^−9^)^^16,800,000^) of not experiencing a substitution along the human branch. Therefore, we conducted the ABC parameter estimation on a set of 10^5^ simulations where we incorporated a 2% rate of random ancestral allele misidentification. Though this binary model of back mutation is highly simplistic (e.g., it does not take into account mutations to another base pair type or trinucleotide context), we found it had only minimal impact on our 95% CIs compared with the same number of simulations that did not incorporate some ancestral state misidentification error (**Table S5**). This suggests that our divergence time estimates may be only slightly underestimated by not accounting for this error.

To investigate the effect of imposing a model of instantaneous speciation rather than bifurcating species divergence on our parameter inference, we also modeled the five gibbon species assuming the best sequence phylogeny from Carbone et al. (in review) and that suggested by our ABC model choice analysis, ((((SSY,HLE)NLE)(HPI,HMO)) (**Table S6**, **Fig. 1b, Fig. S11, Fig. S12**). The seven common median θ values (five extant population values as well θ_T1_ and θ_anc_) were largely concordant, while the 95% CI for θ_T1_ and θ_T2_ were broad and uninformative. Consistent with the rapid speciation hypothesis (even when allowing bifurcating speciation), τ_2+_ τ_3+_ τ_4_ was roughly equivalent to τ_2_ for the instantaneous speciation model, with τ_3_ and τ_4_ being an order of magnitude smaller (i.e., very short internal branch lengths). We also applied the 1 kb data to the method of Gronau et al.(GRONAU *et al.* 2011) as this approach is based on a similar model (i.e., the coalescent with population divergence) as our bifurcating ABC analysis; however, it is more powerful for parameter estimation as it is based on a true likelihood function rather than an approximation (although it does not currently incorporate sequence error rates). While the implementation for estimating divergence times is slightly different (e.g., our ABC approach uses time intervals between divergence events rather than absolute divergence times from the present), the results are very similar: very short internal branch lengths among gibbon genera and a total gibbon genera divergence time of ∼5–6 Mya. However, as expected the 95% CI estimated by G-PhoCS were substantially narrower (**Table S1**).

### Allele Sharing and D-statistic analysis

Because of the small sample sizes and large divergence times it is not expected that we would have power to infer gene flow if added as an additional parameter (whether an instantaneous pulse or continuous migration after divergence) in our ABC analysis. It is also difficult to determine how gene flow would be parameterized in our ABC model framework. Although inter-genera hybrids have been observed in captivity, they are almost certainly infertile as a result of the complicated patterns of homology that would disrupt meiotic pairing. Moreover, such matings have never been observed in the wild, even for sympatric species (HIRAI *et al.* 2007). Therefore, it is unlikely that gene flow would continue for long after divergence as is typically modeled using isolation with migration approaches. Of course, this assumption depends on the rate of karyotypic change, which is thought to have occurred relatively soon after divergence and to have contributed to the speciation process (Carbone et al. in review). Thus, accounting for biologically meaningful gene flow would increase the complexity of the model beyond what can likely be reliably inferred using ABC for this data set.

However, a fairly simply measure that can help to infer admixture events (although not necessarily help to reveal mode, timing or extent of admixture) is the D-statistic (DURAND *et al.* 2011). We first examined patterns of allele sharing across the whole genome by tallying the state of each genus at variable sites by a) choosing sites that met certain quality criteria (as determined by our masks) and that were homozygous for the same allele in both individuals from a genus (filt1) b), randomly sampling one allele from the two genotypes from a genus for sites that met the same quality criteria as a) (filt2), or c) randomly sampling one read from both individuals in a genus at a site (filt3) (**Table S7a**). We also repeated this at the species level, using only the highest coverage sample from each species (in this case filt1 reflects homozygous allele sharing) (**Table S7c**). Results were not qualitatively different using these different filtering criteria.

Consistent with our ABC analysis and Wall et al. (WALL *et al.* 2013), SSY and HLE share the largest number of alleles. Interestingly, while NLE and the two *Hylobates* samples share a fairly low number of alleles compared with other pairwise comparisons, they both share more alleles with SSY than HLE. We performed a D-statistic analysis that demonstrated this excess sharing was statistically significant (**Table S7b and d**). Under the assumption that SSY and HLE diverged last among the four genera as indicated in our ABC analysis, such a pattern is consistent with a model involving two independent gene flow events into SSY from both NLE and *Hylobates* after they diverged from HLE. An alternative model that does not invoke post divergence gene flow involves the maintenance of long-term population structure between the ancestors of HLE and the ancestral population giving rise to the other gibbon genera (**Fig 2**). We attempted to incorporate population structure into our ABC framework but found we had almost no power to distinguish between these models, especially given a parameter space consisting of short internal branch lengths as observed in this data set (data not shown).

**Figure 2b:**
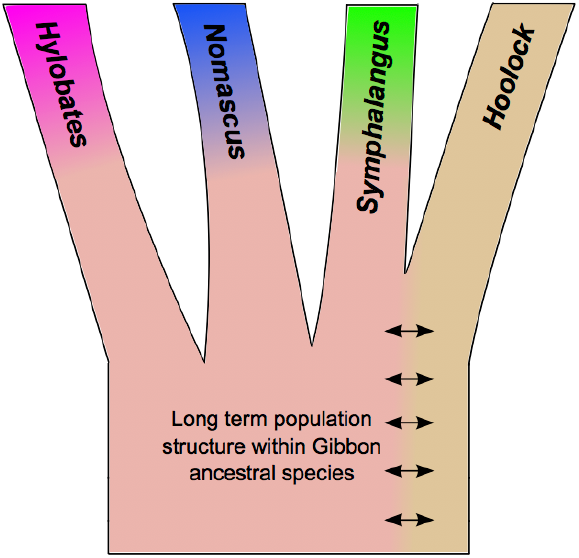
Parameter estimates for a bifurcating speciation model for gibbons genera. µ = 7.5×10^−9^ /site/generation, 10 years per generation. θ_T2_ and θ_T3_ based N_e_ values not to scale.

We also used the D-statistic to examine whether there was any evidence of unbalanced allele sharing between the two Hylobates species. While the D-statistic slightly favored more allele sharing between HMO and the other three genera, the values were generally quite low and the Z-scores were only greater than |2| under filtering scheme 2.

## Discussion

Previous attempts to resolve the phylogenetic relationships among the four gibbon genera based on different genetic systems (karyotypes changes, mtDNA, the Y chromosome, and short autosomal sequences, and ALU repeats) resulted in widely discordant phylogenies. All samples utilized in this study were also analyzed as part of the Gibbon Genome Project (Carbone et al. in review) where the best supported overall consensus tree based on genome-wide sequence divergence was found to be (((SSY, HLE), NLE),(HPI,HMO)). However, all four gibbon genera demonstrated a narrow range for sequence divergence (1.08–1.12%; mean 1.10%). Here, we develop a potentially powerful species tree analysis framework for four taxa that makes use of genome-wide 2^nd^ generation sequencing data and takes into account discordant gene trees, and apply it to the problem of the phylogenetic relationships of the four gibbon genera. Despite our novel methodological approach and the availability of whole genome sequence data, we could not confidently resolve the phylogenetic relationships between *Nomascus*, *Symphalangus*, *Hylobates* and *Hoolock*, although *Symphalangus* and *Hoolock* appear to represent the most recently diverged genera. This result is consistent with the best consensus gene tree identified by Carbone et al (in review) and Wall et al. (WALL *et al.* 2013).

The best topologies are characterized by long external branch lengths and very short internal branch lengths, pointing to a rapid radiation of the four gibbon genera from a large ancestral effective population of ∼10^5^ individuals. This demographic scenario would explain previous observations of genome-wide ILS (Carbone at al. in review) (WALL *et al.* 2013) and discordant phylogenies across smaller datasets. However, we note that an alternative explanation is that the ancestral gibbon population already exhibited structure prior to the divergence of the four gibbon genera.

It is possible that that such a stark restructuring of the gibbon population during this proposed radiation event was driven by some major climatic or geological shift. This is particularly likely as gibbons reside predominantly on the relatively shallow Sunda continental shelf of Southeast Asia. At various times sea level changes and volcanic activity significantly altered the amount of habitable land (i.e., above sea level) in this region. As gibbons live a highly arboreal lifestyle, any reduction or fragmentation of their native forest habitats could have led to extreme genetic isolation between geographically dispersed populations. This, coupled with a rapid evolution of karyotype differences, could have driven the speciation process among these gibbon taxa.

### Uncertainty in timing of the gibbon radiation

It is important to note that associating the timing of speciation with the geological or climatological record is complicated by uncertainty in how we calibrate our estimates of τ (i.e., our choice of mutation rate). A phylogenetic estimate of µ for great apes that is often used is ∼1×10^−9^ per year, an estimate based on calibrating sequence divergence with the fossil record (TAKAHATA and SATTA 1997; NACHMAN and CROWELL 2000). This would place the radiation of gibbon genera within the early Pliocene ∼5 Mya. Interestingly, it has been proposed that the Sunda shelf was largely one land mass up to 5 Mya (OUTLAW and VOELKER 2008), after which sea levels began to rise until ∼3 Mya (CICHON *et al.* 2004) leading to the fragmenting of the region. There is evidence for an increased rate of divergence in other plants and animals during this early Pliocene window (GOROG *et al.* 2004; OUTLAW and VOELKER 2008; AKULA *et al.* 2010; LÓPEZ-GUILLERMO *et al.* 2010) and thus, it is possible that gibbon divergence may have been driven by the same process.

On the other hand, a value of µ = 0.5×10^−9^ per year has recently been estimated using direct observation of trios and quartets in humans (ROACH *et al.* 2010; KONG *et al.* 2012). Scally and Durbin (SCALLY and DURBIN 2012) attempted to reconcile the phylogenetic and direct pedigree estimates with the fossil record (which itself is used to calibrate the phylogenetic estimate) by invoking the hominid slowdown hypothesis. Under this hypothesis, the increased body size of great apes correlates with a decrease in generation time and a reduction in the annual mutation rate after their divergence from Old World Monkeys. Evidence for this comes from evolutionary comparisons of Great Apes to Old World Monkeys (e.g., humans have a 30% slower evolutionary rate as compared to baboons (KIM *et al.* 2006)). While generally bigger than Old World Monkeys, the largest gibbons, from the genus *Symphalangus,* are approximately half the size of the smallest great ape, *Pan paniscus*. Thus, given that gibbons have smaller body sizes (and shorter generation times) than other apes, it is not clear to what extent the hominid slowdown hypothesis would apply.

Decreasing the mutation rate would lead to a Late Miocene speciation time of up to ∼10 Mya, thus encompassing previous estimates of divergence at ∼6–8 Mya based on mtDNA (CHAN *et al.* 2010; VAN NGOC *et al.* 2010; MATSUDAIRA and ISHIDA 2010). However, fossil calibration-based estimates such as used in these studies are subject to their own biases (LUKOSCHEK *et al.* 2012), while estimates of demography from a single locus (especially a non-recombining region of the genome, no matter how well resolved the gene tree) are subject to large evolutionary stochasticity (ROSENBERG and NORDBORG 2002). It is noteworthy that the Y chromosome estimate differs from the mtDNA estimate substantially (5 and 9 Mya respectively) despite application of the same calibration procedures (CHAN *et al.* 2012).

Our results do appear to rule out the hypothesis of Chivers (CHIVERS 1977), which suggests a Late Pleistocene divergence of gibbon genera. Despite this, the constant formation and destruction of land bridges during the Pleistocene that drives the Pleistocene pump hypothesis (GOROG *et al.* 2004; AKULA *et al.* 2010) may have contributed to divergence of the several species within each gibbon genus (for example the *pileatus/moloch* split we observe ∼1.6 Mya). Though the exact numbers are the subject of some debate, it is generally accepted that there are at least 7, 6 and 2 different *Hylobates, Nomascus and Hoolock* species respectively. Movement during these periods likely explains the current distribution of *Hylobates* species both on the mainland and the islands of Sumatra, Borneo, and Java, especially when one considers that gibbons probably cannot swim. Today gibbon species are largely isolated from each other by rivers. Further whole genome sequencing of multiple individuals from additional species, along with the application of powerful genomic methods to infer gene flow or admixture between species, will provide invaluable information for inferring the relationships among gibbon species across Southeast Asia. In addition, while it is well recognized that land bridges certainly formed during the Pleistocene, there is still great uncertainty as to whether these would have involved forest canopy or more savannah-like vegetation (BIRD *et al.* 2005). Analysis of patterns of historic gene flow among the tree dwelling gibbons may help shed light on this process. Recent work using small amounts of autosomal sequence data (∼11 kb) has already found evidence of asymmetrical gene flow between *Hylobates* species currently located on different islands (CHAN *et al.* 2013) while a basic D-statistic analysis in this paper also hinted at the possibility of introgression between genera after divergence.

## Challenges in the use of whole genome sequence data for estimating demographic parameters

Despite the fact that we generated whole genome sequences, it is important to appreciate that the explicit ABC modeling performed here utilized only a small amount of the total available data. A Pairwise Sequentially Markovian Coalescent (PSMC) analysis presented in Carbone et al. (in review) takes a different approach to utilizing genome scale sequence data. By incorporating patterns of genetic diversity across individual genome sequences, important insights can be gained into changing N_e_. To summarize these findings in the context the ABC demographic analysis presented here, both the NLE and HMO populations show major fluctuations in population size during the timeframe after gibbon genera diverged, when Pleistocene geological and climate shifts were taking place.

However, to fully exploit whole genome data for demographic inference using coalescent methods it is vital to construct genetic maps in gibbons, preferably separately for each genus, such that recombination can be appropriately incorporated into the analysis. In addition, despite applying a correction factor in our analysis, reference bias towards *Nomascus* genomes was evident in our data, and it is likely that even more reference bias exists than we actually observe due to the variable karyotypes across genera. It seems unlikely that further large-scale Sanger sequencing will be used to link up scaffolds or generate reference genomes for the other three non-*Nomascus* genera, while short-read Illumina data will have limited power for addressing these aspects. However, the application of new sequencing technologies with long reads such as the PacBio (ENGLISH *et al.* 2012) and nanopore (SCHNEIDER and DEKKER 2012) technologies may provide useful and relatively low cost alternatives to assemble more robust reference genomes. This should lead to more powerful demographic and evolutionary analyses of gibbons in the future.

### Using ABC in Phylogenetics

There is currently one published generalized ABC phylogenetic approach (ST-ABC) (FAN and KUBATKO 2011). This method relies on having accurately phased sequence as it treats frequencies of gene tree topologies across loci as the data rather than summary statistics, and has only been tested and applied to relatively small datasets. However, it has also been questioned whether ST-ABC can accurately approximate the posterior distribution, as it relies on expectations of the distribution of gene trees rather than random simulations that incorporate sampling variability (BUZBAS 2012). Our ABC approach does not have these limitations. While it could not reliably infer the gibbon genera topology with any confidence because of the extremely short internal branch lengths (as predicted by our power analyses), simulations suggest our ABC approach has substantial power to infer the correct species topology for four taxa in most reasonable cases.

However, it is important to appreciate that the framework applied here is tailored for this particular dataset involving unphased genome-wide data from a few individuals per taxa that diverged within the last 10 million years or so. How it would scale up with regard to speed with increasing numbers of samples, and how much power would be lost with fewer loci requires further investigation. It is possible that adding variance in the number of shared sites across loci as a summary statistic may prove useful in this case. In addition, increasing the number of taxa considered (even by one) could prove problematic due to a rapid increase in the parameter space (i.e., a large increase in the number of possible topologies) and an increase in the numbers of summary statistics needed to capture the phylogenetic structure (i.e., the potential impact of the “curse of dimensionality”). Combining more efficient ways of traversing tree space (BRYANT *et al.* 2012) (WEGMANN *et al.* 2009) may help with regard to the former issue, while choosing a more efficient set of summary statistics (e.g., via PLS) may improve the latter; however, there are still likely to be limits to how well the data can be summarized in just a few summary statistics for large phylogenies. Another potential issue of this approach that would place limits on the possible time depth for the phylogeny considered is the assumption of an infinite sites mutation model. It would be trivial to incorporate more complex substitution models, although this would also increase the computational burden.

With these improvements in mind, the ABC-family of methods have the potential to provide a useful and flexible phylogenetic tool that balances the need to incorporate large genomic datasets while taking into account gene tree uncertainty and variation in a coalescent framework. Genomic data is being generated at a rapid pace for a diverse set of species and it clear is that phylogenetic methods are required than can accommodate such data. ABC provides one approach to do this.

## Acknowledgments

Support for this work was provided by the National Institutes of Health to J.D.W and M.F.H (R01_HG005226). We thank Ryan Sprissler and the University of Arizona Genetics Core for assistance with sequencing and Ryan Gutenkunst for computing resources.

---

**Supplementary Figure 1: Cartoon showing the distribution of genic (200bp) and non-genic (1kb) loci identified for phylogenetic analysis of gibbons**. Not to scale.

**Supplementary Figure 2: Distribution of coverage in the 8 gibbon samples.**

**Supplementary Figure 3: PCA of all 8 gibbon samples based on high quality genotypes.** A) PCA 1v2 for all SNPs, B) PCA 3v4 for all SNPs, C) PCA 1v2 for random 1% of SNPs, D) PCA 3v4 for random 1% of SNPs.

**Supplementary Figure 4: Relative posterior probabilities as assessed by our ABC framework for 10,000 random simulated topologies (from a total of 15 possible topologies for 4 genera) using the Logistic Regression (LR) and Direct (DR) methods.** Simulations are ordered from highest to lowest LR posterior probabilities.

**Supplementary Figure 5: Posterior probabilities of the true model (either a asymmetric or symmetric tree from a total of 15 possible models or topologies) as assessed by our ABC framework for a specific framework of demographic scenarios using the Direct (DR) and Logistic Regression (LR) methods.** T_anc_ equals the total height of the tree. Conversion of height from substitutions per site to years is based on a mutation rate of 1×10–9 per year. For θ = mixed, see Supplementary text for exact parameterization.

**Supplementary Figure 6: (A) PCA of corrected simulated and observed (red) summary statistics for (A) non-genic loci and (B) genic loci.**

**Supplementary Figure 7: Posterior distributions for the instantaneous speciation model for gibbon genera.**

**Supplementary Figure 8: RMSE of PLS components for the instantaneous speciation model for gibbon genera.**

**Supplementary Figure 9: Posterior distributions for the instantaneous speciation model for gibbon genera using flat priors for for τ.**

**Supplementary Figure 10: RMSE of PLS components for the instantaneous speciation model for gibbon genera using flat priors for for τ.**

**Supplementary Figure 11: Posterior distributions for a bifurcating speciation model for gibbons genera.**

**Supplementary Figure 12: RMSE of PLS components for the bifurcating speciation model for gibbon genera.**

**Supplementary Figure 13: Net missed singletons per base, as a function of read depth.** The net missed singletons per base is computed on the intersection of the exome-capture and the whole genome data, and is given as a function of the read-depth in the whole genome data. Due to sampling variance, there is modest variance in this function, but nevertheless error rates initially are high, then decrease, and then slightly increase again.

**Supplementary Figure 14: Genotyping error rates.** For sites not called as singletons in the whole genome data, the error rate (defined as the probability of a miscall), is given as a function of the read-depth. Even after filtering CNV regions, error rates increase at high read-depth.

**Supplementary Figure 15: Example model setup for an asymmetric phylogeny.**

**Supplementary Figure 16: Example model setup for a symmetric phylogeny.**

